# Sustained bias of spatial attention in a 3T MRI scanner

**DOI:** 10.1101/2024.03.26.586826

**Authors:** Stefan Smaczny, Leonie Behle, Sara Kuppe, Hans-Otto Karnath, Axel Lindner

**Author notes:** Shared Correspondence (;)., *<u>Addresses for correspondence:</u>* Axel Lindner, Center of Neurology, University of Tübingen, Hoppe Seyler Str. 3, D-72076 Tübingen, Germany, Hans-Otto Karnath, Center of Neurology, University of Tübingen, Hoppe Seyler Str. 3, D-72076 Tübingen, Germany. These authors contributed equally to the paper (Stefan Smaczny, Leonie Behle).

## Abstract

The static magnetic field of MRI scanners induces magneto-hydrodynamic stimulation of the vestibular organ (MVS) in the inner ear. This not only causes a horizontal vestibular nystagmus but also induces a horizontal bias in spatial attention. In this study, we aimed to determine the time course of MVS-induced biases in both VOR and spatial attention inside a 3T MRI-scanner as well as their respective aftereffects after participants left the scanner. Eye movements and overt spatial attention in a visual search task were assessed in healthy volunteers before, during and after a one-hour MVS period. All participants exhibited a VOR inside the scanner, which declined over time but never vanished completely. Importantly, there was also an MVS-induced horizontal bias in spatial attention and exploration, which persisted throughout the entire hour within the scanner. Upon exiting the scanner, we observed aftereffects in the opposite direction manifested in both the VOR and in spatial attention, lasting for about 6 minutes. Sustained MVS effects on spatial attention have important implications for the design and interpretation of fMRI-studies and for the development of therapeutic interventions counteracting spatial neglect.

## Introduction

Strong magnetic fields such as those permanently present in magnetic resonance imaging (MRI) scanners stimulate the vestibular organ in the inner ear and induce a persistent nystagmus consisting of a slow vestibular ocular reflex (VOR) followed by quick resetting saccades in the opposite direction. This phenomenon is due to a magneto-hydrodynamic vestibular stimulation (MVS) and has been demonstrated in various studies using MRI scanners with different field strengths and exposure durations (Roberts et al., 2011; Ward et al., 2019). MVS is caused by an interaction between the static magnetic field of the scanner and ionic currents in the endolymph fluid of the subject’s labyrinth, producing a Lorentz force (Roberts et al., 2011). Only recently, Lindner et al. (2021) showed in healthy individuals that MVS not only causes a vestibular nystagmus but also affects spatial attention and, more specifically, spatial orientation and exploration behavior: In a visual search task inside a 3T MRI scanner, the healthy subjects’ spatial attention was – like the VOR - significantly shifted towards the right.

These latter results resemble the well-known effects of caloric vestibular stimulation (CVS) on spatial orientation and exploration in healthy subjects. CVS likewise not only induces a VOR but also a bias of subjects’ spatial attention and exploration: when exploring their surroundings for possible targets, subjects’ eye movements are no longer symmetrically distributed in the horizontal dimension but biased towards one side of the body’s midsagittal plane (Holé et al., 2020; Karnath et al., 1996; Karnath, 1994). However, the effect of CVS lasts only a few minutes, whereas the physiological effect of MVS (as estimated through the VOR) remains throughout the entire time while the subject is inside the scanner (Jareonsettasin et al., 2016; Roberts et al., 2011). This has been demonstrated for a duration of 90 minutes inside a 7T scanner in healthy subjects, in whom nystagmus initially diminished but did not extinguish (Jareonsettasin et al., 2016). After such prolonged MVS there was, in addition, a post-stimulatory aftereffect that was present immediately after exiting the scanner. This aftereffect was characterized by slow phases of the nystagmus in the opposite direction as compared to inside the magnet. The aforementioned effects can be interpreted as the result of a VOR set-point adaption process due to prolonged exposure to MVS (Jareonsettasin et al., 2016).

First evidence at 3T demonstrates that the VOR is likewise sustained for periods of about 20 minutes (Go et al., 2022). Still, it might be possible that there is a complete cessation of the VOR at lower field strengths for longer MVS exposure times. In contrast to the initial findings on the VOR (Go et al., 2022), it is currently still completely unknown whether the behavioral shift in spatial orientation and exploration is likewise sustained during prolonged MVS. If present, these sustained effects of MVS could have major implications in at least two domains. First, they bear the potential to ameliorate pathological spatial attention biases in stroke patients suffering from spatial neglect. Spatial neglect is a lateralized spatial attention disorder that occurs predominantly after right hemisphere lesions (Karnath & Rorden, 2012). It is known that this disorder can be markedly ameliorated - even fully compensated - through CVS (Karnath et al., 1996; Karnath, 1994; Rubens, 1985; Vallar et al., 1995) as well as through MVS (Karnath et al., 2022). In contrast to CVS, however, MVS-induced corrective shifts of spatial attention and exploration could serve as a much more effective therapeutic tool if the behavioral shift would sustain during prolonged MVS. Second, any sustained effects of MVS on both spatial attention and the VOR need to be considered when designing and interpreting any fMRI studies in healthy subjects (Boegle et al., 2016; Lindner et al., 2021).

In the present study, we thus investigated the time course of MVS effects during and after a one hour exposure to a 3T magnetic field, namely on both the VOR (as a proxy for the effectiveness of the vestibular stimulation) and on spatial attention and exploration. We expected to replicate an initial MVS-induced VOR with slow phases of the nystagmus to the right, due to the magnetic field direction of our scanner (Boegle et al., 2016; Jareonsettasin et al., 2016; Roberts et al., 2011), as well as a rightward shift of attention in visual search (Lindner et al., 2021). We asked whether or not this spatial attention shift in the visual search task would persist along with the rightward VOR throughout one hour of stimulation in the 3T scanner, thereby resembling the sustained time course of the VOR during MVS at 7T (Jareonsettasin et al., 2016; Roberts et al., 2011). Moreover, with respect to previous findings of a MVS-induced VOR aftereffect at 7T (Jareonsettasin et al., 2016; Zee et al., 2017), we expected a smaller and gradually diminishing aftereffect in the opposite direction for both the VOR and for spatial attention immediately after subjects leave the MRI bore.

## Methods

### Subjects

Thirteen participants were recruited for the study. The number of subjects resulted from a power analysis using data from the search task collected by Lindner et al. (2021), as this was our main measure of interest (one-tailed tests with alpha = 0.05, a power of 0.9 and an expected effect size of Cohen’s *d* = 0.88). The eye-tracking data of one subject could not be analyzed due to artifacts introduced by the scanner-compatible glasses. Hence, only 12 datasets could be analyzed (N=12, 3 males, average age was 24 ± 2.9 years, 1 subject was left-handed). All subjects were screened for MRI compatibility to exclude those with general MRI restrictions, neurological or psychiatric histology, claustrophobia, or fear of the dark. The experiment lasted about 2 hours and participants received compensation. All gave their signed, informed consent according to the institutional ethics board guidelines prior to the experiment. The study followed the Declaration of Helsinki and was approved by the Ethical Committee at the Medical Faculty of Tübingen University.

### General Procedure

Three different tasks (eye calibration, central fixation in darkness to assess the VOR, and visual search; compare below) were performed by participants in overall nine runs outside and inside the scanner. An overview of the time course of the experiment is depicted in Figure 1. One run with all three tasks lasted 5.7 min on average. We conducted one run outside the scanner at the beginning (Pre 1). Pre 1 served to provide us with baseline estimates for the VOR and visual search in the absence of MVS. This “pre outside phase” was followed by the “inside phase” with three runs inside the MRI bore (In 1-3): In 1 was carried out immediately after subjects had entered the bore, In 2 was performed after about 27 minutes since subjects had entered the bore, and In 3 was conducted after about 55 minutes and immediately before subjects left the bore. These runs allowed us to examine whether we could induce the expected MVS-effects (comparing respective measures for In 1 vs. Pre 1) and to probe how they develop over time (comparing In 1-3). Once subjects had left the scanner, they completed five additional runs in direct succession (Post 1-5) to assess any MVS-induced after-effects during this “post outside phase”. To prevent fatigue or falling asleep while lying inside the scanner, radio documentaries were presented between the sessions inside the scanner (approximately 20 minutes each). The documentaries were pseudorandomized across participants. To prevent any effects of the content on our experiments, such as spoken directions, each documentary was played to a maximum of three different participants.

**Figure 1.**
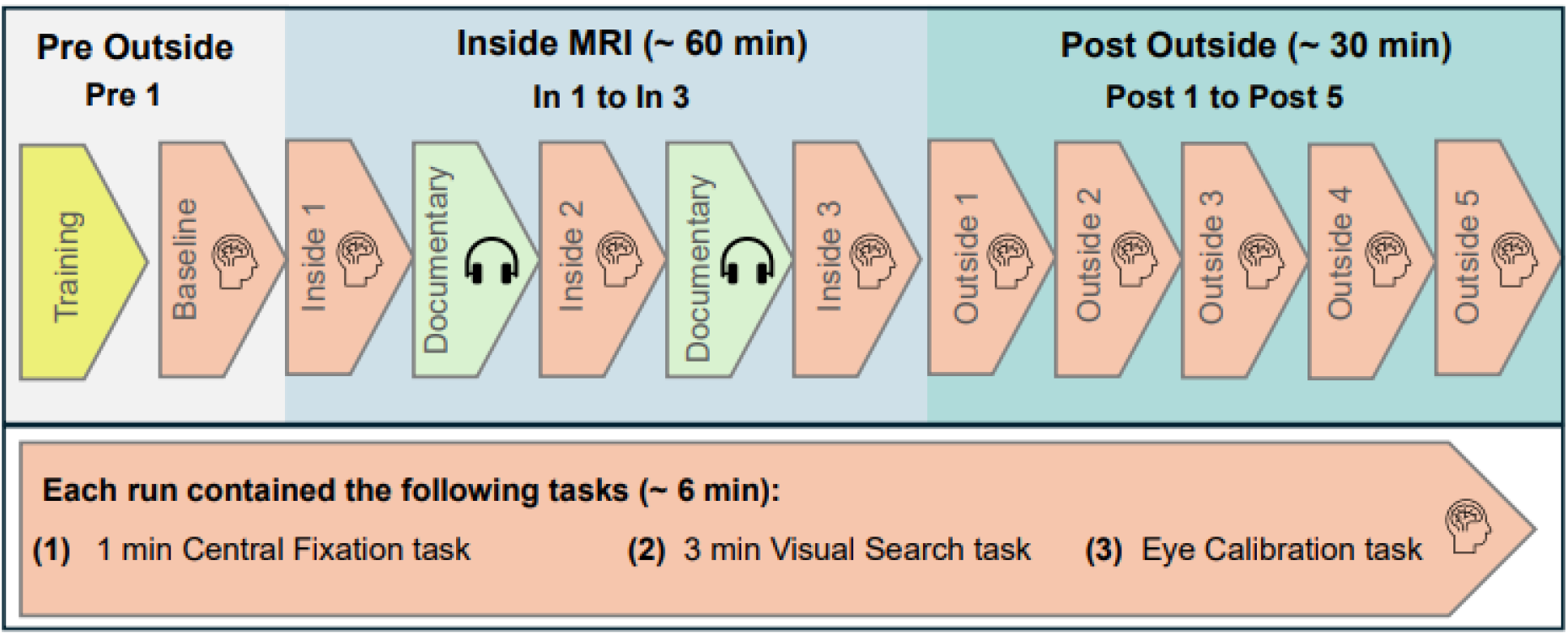
Overview over the time course of the experiment.

First, subjects performed a *central fixation task*, in which a central light stimulus was presented for 5 seconds. Participants were instructed to fixate on the light and, after the light disappeared, keep fixating this point for one minute. Eye position was measured via eye tracking. This task was carried out to quantify participants’ VOR. We used the median de-saccaded horizontal eye velocity as a measure for the VOR in each individual. Second, participants had to conduct a *visual search task*: participants had to scan their visual field for about three minutes (172 s) to find small LED lights. During this time course, six LED lights were presented at random times for five seconds each, with the voltage doubling each second from 0.1 to 1.6 V, making it more likely for participants to find the target. The presentation of these stimuli only served to maintain the subjects’ motivation to search for possible targets. Our interest was the spatial distribution of subjects’ overt spatial attention, as assessed by their exploratory scan path in the absence of any visual targets. The final target was always presented centrally for two seconds at maximum intensity of 5 V for calibration purposes. The x/y positions of the search targets were -6°/5°; 6°/5°; -6°/-5°; 6°/-5°; -12°/-1°; 12°/1° and 0°/0° (values indicating visual angle in right/upward direction for positive values and vice versa). For the seven inter-target-intervals (ITIs), subjects were in complete darkness. The duration of the ITIs varied (5 to 35 s in 5 s steps). Both the target order as well as the ITIs were pseudorandomized across the multiple search tasks of each participant and between participants. In addition to fixation, subjects were asked to press a button on an MRI-compatible response pad during the search task whenever they found a target. We used a 5-Button Diamond Response Pad (Current Designs). Note that before starting Pre 1, subjects completed a training for this search task with four targets and 5 ITIs (5-25 s). Finally, participants underwent an *eye-calibration task* in which five LED lights were presented at maximum voltage for 2 seconds each. The x/y positions of the eye-calibration targets were 0°/0°; - 6°/5°; 6°/5°; - 6°/-5°; 6°/- 5°, always shown in this order and this cycle was presented twice, with an overall duration of 20 s. Note that for the post outside phase, this calibration task was performed only at the end of every other run, namely Post 1, Post 3 and Post 5, respectively.

For our experiment, we used a 3T Siemens MAGNETOM Prisma MRI Scanner. No radio frequency or gradient field were applied, as no imaging was performed, and only the static magnetic field was present. Participants entered the MRI head-first and the magnetic field vector pointed from subject’s toes to their head (see Lindner et al., 2021). The experiment was implemented in MATLAB using a WIN 10 laptop PC with custom MATLAB scripts (R2015b 32 bit, MathWorks) in combination with Cogent 2000 and Cogent Graphics (by FIL, ICN, LON at the Wellcome Department of Imaging Neuroscience, University College London) and the MATLAB Support Package for Arduino. The light stimuli were generated with an Arduino UNO R3-compatible microcontroller (Funduino UNI R3) that controlled eight red LED lights (L-513HD; Dropping resistor: 48 kΩ). Eight glass fibers led from the microcontroller into the scanner room and into a black custom-made board at fixed locations. Each glass fiber was connected to one LED on the board to generate one stimulus at a certain location. The board was mounted behind the subject’s head, where subjects could see it (and thus the stimuli) using a mirror on the head coil. Intensity of the LEDs could be adjusted for six LEDs between 0 and 5 V using the Arduino’s pulse code modulated (PCM) analog voltage output. The remaining two LEDs could only be switched on (at maximum intensity of 5 V) or off, although only one of them (at the position 0°/0°) was used. The light stimuli were at a viewing distance of 1.1 m with the glass fiber diameter being 1 mm (0.05° visual angle), which again was equal to Lindner et al. (2021). This distance was chosen based on the findings of (Owens & Leibowitz, 1980), who found that the mean vergence of the eyes in the dark is about 1.1 m. Thereby we tried to reduce any influence of vergence on our monocular eye-tracking data in the presence/absence of light.

Subjects were in complete darkness throughout the experiment, apart from the small, red LED light stimuli. This was realized by attaching a thick, black blanket to the MRI scanner that fell onto the subjects and blocked most light. Additionally, we removed any other light source in the scanner room by covering them with black cardboard. At every new position of the scanner table we ensured that participants could not see any external light.

The head of subjects was pitched back using cushions to achieve a roughly -30° angle between the vertical axis and Reid’s plane (line between the infraorbital margin of the orbita to the upper margin of the external auditory meatus). This was done to maximize the VOR effect, based on data by Roberts et al. (2011) and Boegle et al. (2016), who analyzed the horizontal slow phase velocities depending on head pitch angle. Inside the coil, smaller cushions were placed on both sides of the head to fixate the subject’s head and prevent head movement, especially to the sides.

### Eye Tracking Setup

For eye tracking, a second WIN 10 PC was used. Eye tracking was executed via an MR-compatible camera with integrated infrared LED illumination (MRC Systems; Model: 12M-I IR-LED) at 50 Hz sampling rate. The camera was mounted onto the head coil and monitored the subjects’ right eye via dark pupil tracking during execution of the tasks. To obtain the uncalibrated eye-position data, the eye-camera video was digitized using the ViewPoint Monocular Integrator System and the ViewPoint Software (Arrington Research, software version 2.8.3.437). The WIN 10 laptop PC for the experiment had remote control through the ViewPoint Ethernet-Client of the second WIN PC where the eye tracking was running.

### Eye Movement Analyses

Eye data were analyzed after the experiment using custom-written scripts in MATLAB R2017b (MathWorks). For this analysis, eye-position samples were filtered using a second-order 10 Hz digital low-pass filter. Based on the calibration task described earlier, a five-point calibration was performed and applied to the eye movement recordings from the central fixation task and from the visual search task. This was done separately for most runs: To save time while measuring the after-effect, there were calibrations done at the end of Post 1, Post 3 and Post 5. The calibration after Post 1 was used for Post 1 and Post 2, while the calibration after Post 3 was use for Post 3 and Post 4. Furthermore, an additional compensation for eye position offsets was performed in the central fixation task and the visual search task: Although we immobilized the head of subjects with cushions, even tiny head movement across tasks could have had a substantial influence on our eye-tracking data, thus an additional offset-compensation was applied for each task individually. For the central fixation task, the initial central light stimulus was used to correct any offset, while in the visual search task, the visible search targets themselves guided offset compensation (also compare Lindner et al., 2021).

To determine saccades, an absolute eye-velocity threshold of 15°/s was used, with saccade-onset being defined as the first sample after threshold-crossing. Note that eye velocity was calculated based on two-point differentiation of our eye position data. Accordingly, saccade-offset was defined as the first sample of the eye velocity dropping below the threshold. Blink artifacts were excluded from analyses. For the calculation of the horizontal VOR (slow phase) velocity, the time periods during saccades (from on-to offset) were treated as missing values.

For the search-task data analysis, we focused on the distribution of horizontal saccade endpoints characterizing subjects’ visual scan path (also compare Lindner et al., 2021). Time periods with a search target present were removed and only the ITIs were used (140s). In addition, a 5-second interval was excluded after each target presentation to avoid any carry-over effects due to prior target-fixation, leaving a 110-second time period for saccade endpoint analysis.

### Statistical Analyses

For each participant, the mean horizontal saccade endpoints from the search task (reflecting the center of visual search) and the median of the de-saccaded horizontal eye velocity from the central fixation task (as a measure of the VOR) were calculated in each individual and for each run. All values were normalized by subtracting their respective baseline measure (Pre 1). Across subjects the mean baseline value for the horizontal VOR amounted to 0.32°/s +/- 0.41°/s standard deviation (SD), while the mean horizontal center of visual search was 4.45° +/- 2.68° SD. Both baseline measures well compare to those obtained in a previous study of our laboratory (Lindner et al., 2021). All normalized means were normally distributed (Shapiro Wilk Test, p’s > .01). For both the saccade endpoint and the VOR means we first performed a manipulation check to test whether In 1 and Post 1 differed from the baseline (Pre 1), representing the main effect of MVS and its aftereffect, respectively (one-tailed t-tests). Then we calculated two separate one-way repeated measures ANOVAs. One examined the difference between the inside runs (In 1, In 2, In 3) to ascertain time differences in the effect of MVS on VOR and search behaviour while inside the scanner. The other repeated measures ANOVA was to investigate the temporal course of the after-effect (all outside runs). To examine when the aftereffect wore off, we examined all further outside runs individually with the baseline using one-tailed t-tests. These tests were Bonferroni-corrected for multiple comparisons. The same procedure was also applied to probe whether MVS effects inside the scanner were maintained and thus differed from baseline.

## Results

### Vestibulo-Ocular Reflex

Due to the magnetic field direction of our MRI scanner (magnetic field vector pointing from subjects’ toe to head), nystagmus was expected to develop with the slow phases to the right and resetting saccades to the left inside the scanner. When looking at the individual de-saccaded horizontal eye velocity for In 1, all subjects showed a relative increase in velocity towards the right inside the scanner compared to baseline (In 1: M = 1.32 °/s ± 0.89 °/s SD; see Figure 2). Our subjects completed the tasks inside the scanner for two more times (in the middle and at the end of the one-hour inside phase). Here, the VOR effect was maintained but clearly decreased between In 1 and In 2 (In 2: M = 0.59 °/s ± 0.55 °/s SD) but barely changed from In 2 to In 3 (In 3: M = 0.54 °/s ± 0.51 °/s SD).

**Figure 2.**
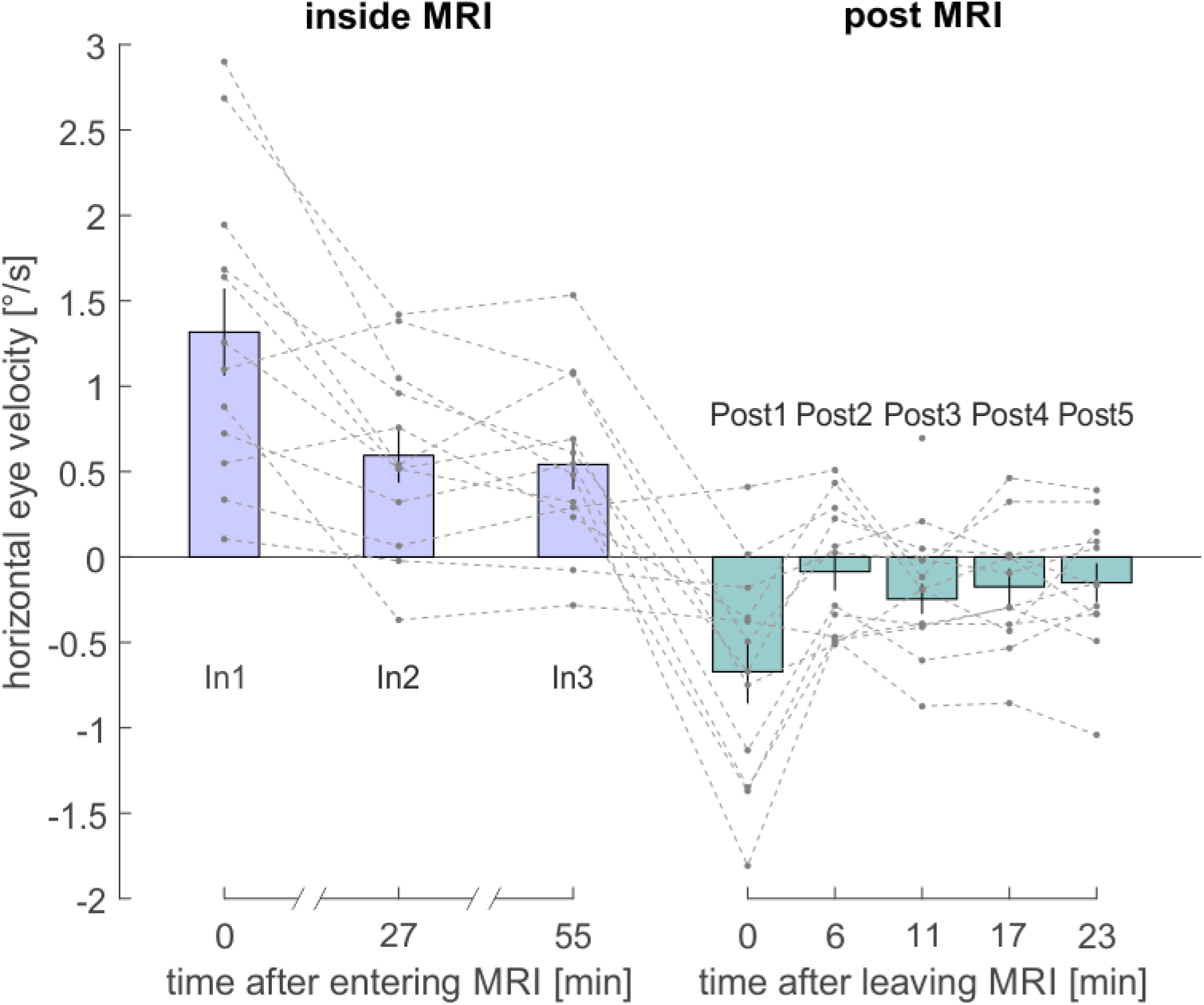
De-saccaded horizontal eye velocity (VOR) in °/s during each run relative to the baseline (Pre 1). Dotted lines show data for each participant. Violet bars show the mean VOR for each run over all subjects inside the MRI bore, turquoise bars show the mean VOR post outside of the MRI bore (Post 1-5). Black error bars indicate the standard error. Measurements of VOR took place right after the subject entered the scanner (In 1) and after 27 minutes (In 2) and after 55 minutes (In 3) lying in the scanner. Further measurements of VOR were performed right after participants were removed from the scanner (Post 1), and repeated every 5.7 minutes (Post 2 to Post 5). In all figures, *time* refers to the average onset of individual runs, rounded to minutes.

We performed a manipulation check to test whether the main effect of stimulation on VOR was evident for In 1. A one-sample t-test comparing In 1 with zero was significant, t(11) = 5.15, p < .001, d = 1.49. We then performed a one-way repeated measures ANOVA, comparing the inside runs with one another to examine whether the effect differed between inside runs. Mauchly’s test indicated that the assumption of sphericity had been violated, *χ2*(2) = 6.18, *p* < .05. Therefore, Greenhouse-Geisser corrected tests are reported (*ε* = 0.68). The ANOVA was significant, *F*(1.37, 15.06) = 12.67, *p* = .001, *η2* = .54. Planned contrasts revealed both a significant linear trend over time, *F*(1,22) = 14.57, *p* = .003 as well as a quadratic trend, *F*(1,22) = 8.31, *p* = .015. The results demonstrate that the VOR during the three measurements inside the scanner decayed in a non-linear fashion but, nevertheless, was sustained throughout the stimulation and was significantly different from baseline: The VOR in In 2 was significantly larger than the baseline, one-tailed, one-sample t-test, *t*(11) = 3.74, *p* = .002, d = 1.08. The same was true for In 3, *t*(11) = 3.69, *p* = .002, d = 1.06.

We next examined whether there was a significant aftereffect right after leaving the MRI bore. A one-sample t-test shows that Post 1 was significantly smaller than the baseline (Pre 1), suggesting the presence of an aftereffect, *t*(11) = -3.62, *p* = .002, d = -1.04. To examine how the aftereffect changed over time, we performed a one-way repeated measures ANOVA, comparing the post outside runs with one another. Mauchly’s test indicated that the assumption of sphericity had been met (χ2(9) =11.43, p > .05. The ANOVA was significant, *F*(1,22) = 11.34, *p* = .001, *η2* = .51. Planned contrasts suggest a significant linear trend over time, *F*(1,44) = 17.95, *p* = .001, indicating that VOR linearly decreased from a leftward bias back to normal. Moreover, a significant quadratic trend was found, *F*(1,22) = 7.86, *p* = 0.017. To examine when the aftereffect wore off, we examined all further post outside runs individually with the baseline using one sample t-tests. After the initial aftereffect in Post 1, none of the further outside runs differed significantly from the baseline (all p’s > .05).

### Visual Search

Due to the magnetic field direction of our MRI scanner, we expected visual search as measured by a subject’s mean horizontal position of saccade endpoints to shift rightward inside the scanner while no target was present. Figure 3 shows an example of some of the measurements for a single participant, while Figure 4 provides an overview of the entire data set for all measurements and all participants.

**Figure 3.**
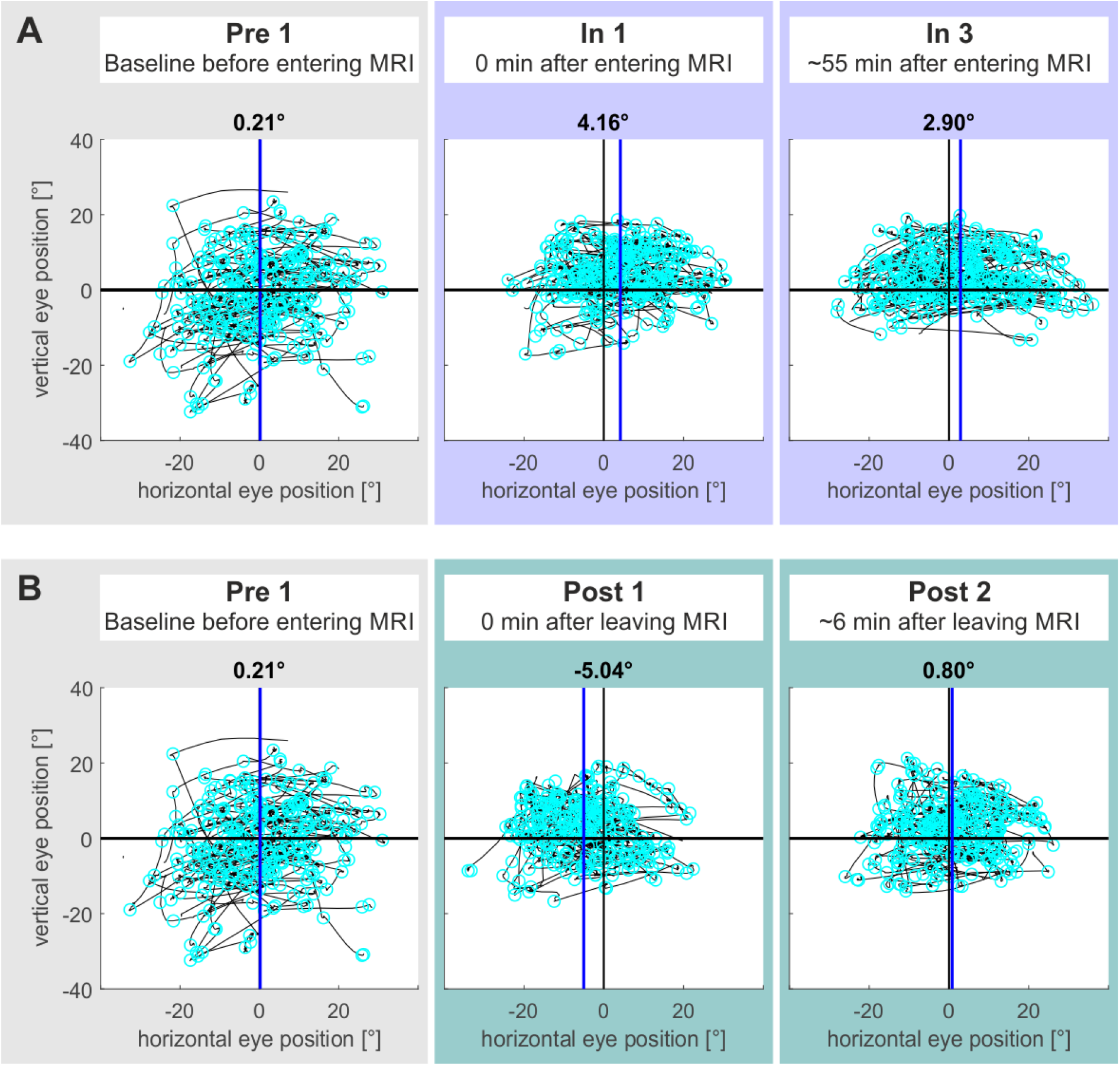
Exemplary data for the visual search task from one exemplary participant for some of the measurements. Black lines show the 2D visual scan path in degrees visual angle during the three-minute search task. Cyan circles depict saccade endpoints. The blue vertical lines show the mean of saccade endpoints. **A** depicts an MVS-induced sustained shift of mean visual search inside the scanner that persisted from the beginning (In 1) until the end (In 3) of the ∼60 minutes inside phase as compared to baseline (Pre 1). **B** depicts an MVS-induced aftereffect in visual search in the opposite direction (Post 1 vs. Pre 1), which already vanished after about 6 minutes (Post 2).

**Figure 4.**
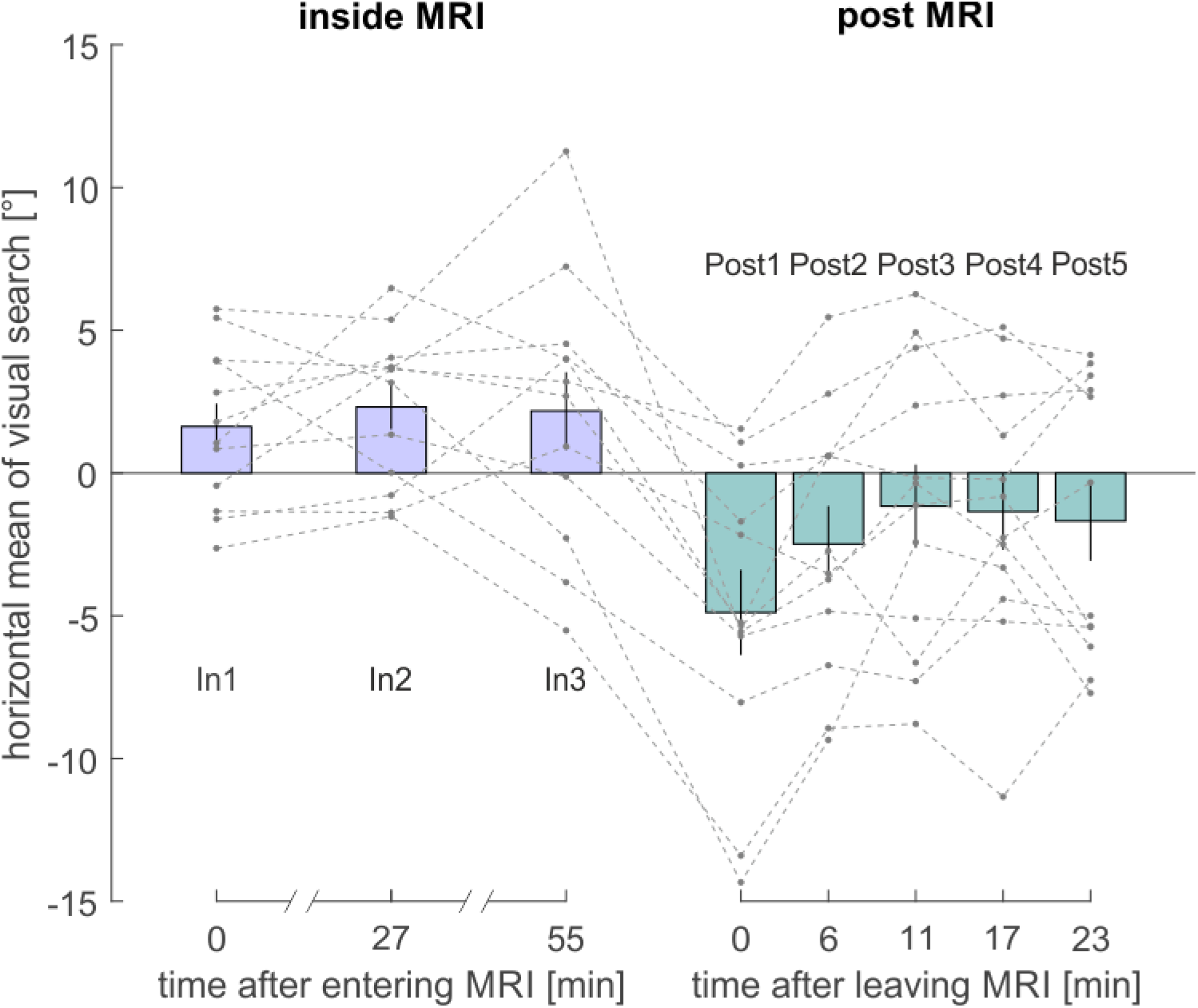
Mean horizontal position of saccade endpoints in ° during the search task relative to the baseline (Pre 1). Dotted lines show data for each participant. Violet and turquoise bars show the mean horizontal eye position over all subjects for inside and post outside MRI, respectively. Black error bars indicate the standard error. Measurements of the horizontal position of visual search took place right after the subject entered the scanner (In 1) and after 27 (In 2) and after 55 minutes (In 3) lying inside the scanner. Further measurements of the horizontal position of visual search were performed right after participants were removed from the scanner (Post 1), and then again every 5.7 minutes lying outside the scanner (Post 2 to Post 5).

When looking at the mean saccade endpoints for each subject, most participants showed a shift in visual search towards the right inside the scanner by a mean of 1.63° ± 2.80 °/s compared to baseline (Figure 4). Our subjects completed the tasks inside the scanner for two more times (in the middle and at the end of the one-hour inside phase). Here, the rightward bias of the visual search towards the right remained stable over all runs (In 2: M = 2.31 °/s ± 2.70, In 3: M = 2.17 ± 4.71).

To first replicate the presence of MVS on search performance immediately after entering the scanner (cf. Lindner et al., 2021), we tested whether there was a difference between In 1 and the outside baseline on the mean horizontal position of search. Participants showed a significantly rightward shift in their visual search, one sample t-test, *t*(11) = 2.01, *p* = .035, d = 0.58. Mauchly’s test indicated that the assumption of sphericity had been met (χ2(9) = 5.72, p > .05). There was no significant difference between the inside runs, one-way repeated measures ANOVA, *F*(1,22) = 0.23, *p* = .798, resembling the fact that there was no noticeable change in visual search bias over runs across subjects. Accordingly, the linear (*F*(1,22) = 0.16, *p* = 0.694) and quadratic (*F*(1,22) = 0.48, *p* = 0.502) trends were not significant. As we did not observe relevant differences between inside runs, we pooled the data from In 2 and In 3 for each participant individually and calculated a one sample t-test against zero to examine whether the effect of MVS was present after subjects had been in the scanner for about 27 or 55 minutes. There was a significant difference between the pooled values and zero, *t*(11) = 2.30, *p* = .021, d = 0.66. Thus, the effect of MVS was present for the entire time participants spent inside the MRI bore.

As for the VOR, there was also a significant aftereffect in visual search right after subjects left the MRI bore. A one-sample t-test showed that Post 1 was significantly smaller than the baseline (Pre 1), suggesting the presence of an aftereffect with a bias towards the left, *t*(11) = -3.27, *p* = .004, d = -0.94. To examine how the aftereffect changed over time, we performed a one-way repeated measures ANOVA across the post outside runs. Mauchly’s test indicated that the assumption of sphericity had been met (χ2(9) = 15.56, p > .05). The ANOVA was significant, *F*(2.58, 28.36) = 6.68, *p* <.001, *η2* = .38. Planned contrasts suggest a significant linear trend over time, *F*(11,44) = 13.89, *p* = .003, as well as a quadratic trend over time, *F*(11,44) = 6.34, *p* = .029, indicating that visual search decreased non-linearly from a leftward bias back towards the middle. To examine when the aftereffect wore off, we examined all further post outside runs individually with the baseline using one sample t-tests. After the initial aftereffect in Post 1, none of the further out-runs differed significantly from the baseline (all *p*’s > .05). Therefore, the after effect only lasted for approximately 5 minutes.

## Discussion

The present study set out to determine the temporal dynamics of the MVS-induced VOR and the spatial attention bias inside a 3T MRI scanner over a period of one hour. Additionally, we investigated the duration of any resulting aftereffect of the prolonged MVS-exposure on behavioral responses. MVS at 3 Tesla induced a rightward VOR, which declined but never completely subsided throughout the period of one hour inside the scanner. Importantly, MVS also induced a horizontal rightward bias in spatial attention and exploration that was stable over time, emphasizing a robust and sustained effect of MVS on visual search. Finally, upon exiting the scanner, we observed aftereffects in both VOR and spatial exploration, which were in the opposite direction and which were significant within the first 6 minutes of the outside phase, only.

An earlier study by Jareonsettasin and coworkers (2016) already studied the influence of MVS on the VOR at 7 T and for time periods of up to 90 minutes. Our results at 3T are well compatible with their finding of a sustained VOR and a reported ∼300 s time constant of VOR adaptation: Immediately after entering the scanner (In 1), the VOR was most pronounced and exhibited a significant decay to a stable level already at In 2 (i.e. after 27.5 minutes). Importantly, also for our field strength of “only” 3T (in contrast to 7T in their study) was this sustained level of VOR clearly different from baseline. While Go and coworkers (2022) have already demonstrated such sustained VOR at 3T for a scanning time of about 20 minutes, we further show that this effect holds on for a time period of at least up to one hour. Finally, the findings that our VOR after-effect was present only for Post 1 and that it was no longer statistically detectable thereafter (i.e. at Post 2 after 6 minutes, etc.), is again compatible with a respective ∼100 s time constant reported by Jareonsettasin et al. (2016) at 7T. Note, that our approach did not allow us to derive precise time constants as we could not obtain time-continuous measurements of the VOR. This was due to interleaved calibration and visual search tasks that were needed to address a completely novel aspect of our work, namely how MVS affects spatial attention over longer periods of time.

In fact, we observed that the MVS-induced rightward shift in spatial attention was constantly present throughout the overall duration of our 60 minutes inside phase. Different to the VOR, however, this bias did not exhibit any adaptive decline over time. That some adaptation of the MVS-induced bias in spatial attention must have still taken place was evident from the significant after-effect during (and only during) the first measurement interval after subjects had left the scanner (Post 1). Most importantly, however, both MVS-induced VOR and attentional bias were present throughout the total duration of the inside phase in the 3T scanner.

When considering that the duration of typical fMRI experiments is on the same order as our inside phase, questions about the impact of these sustained effects of MVS on behavioral and functional measures do arise. First, others have summarized diverse vestibular influences on cognition (e.g. Ferrè & Haggard, 2020), which could affect behavioral performance during fMRI. Moreover, the fact that MVS influences neural network activity has been elegantly demonstrated before (Boegle et al., 2016, 2017, 2020). Given that vestibular influence on brain activity is also present under natural lighting conditions, i.e. even when the VOR is suppressed (see Go et al., 2022; Lindner et al., 2021; Ward et al., 2017), our present results indicate that during the entire fMRI acquisition time the neuronal activation in the (cortical) projection areas of the vestibular system represent an unresolvable mixture of the signal of interest and the signal evoked by the activation of the vestibular system. Such neural consequences of MVS should be critically considered in any fMRI study in which activations are expected in networks also affected by MVS. Moreover, measures of lateralized activity and behavior might strongly depend on the direction of MVS and on associated biases in spatial attention.

Apart from these challenges, the sustained effects of MVS do − as compared to any other vestibular stimulation technique (Ertl & Boegle, 2019) – present a unique opportunity to study vestibular influences on cognition and brain activity. For example, Jareonsettasin and coworkers (2016) studied adaptation of the VOR in an attempt to understand how the nervous system reduces an unwanted nystagmus that is induced by a “vestibular pathology”, virtually induced through MVS. In other words, they studied VOR adaptation towards a new “set point” at which the head should be considered as stationary in space despite the “pathological” vestibular signaling leading to unwanted nystagmus. Others used MVS to study “vestibular perception” (Mian et al., 2013, 2015, 2016).

Beyond such opportunities for basic research, MVS also promises new possibilities for the effective treatment of vestibular but also other diseases, such as, e.g., spatial neglect following stroke. Stroke patients with spatial neglect act in a way as if the “set point”, which defined the center of their behavior with respect to their body midline, was pathologically shifted to the right (Karnath, 1994b, 2015). At a more abstract level, a set point defines a “resting position”, from which to launch movements, and “set point adaptation” refers to the nervous system’s attempt to continuously maintain stability of a set point despite development and disease (cf. Zee et al., 2017; also compare the aforementioned example of set point adaptation of the VOR). Any therapeutic intervention, which would reduce the pathological bias in patients’ exploratory behavior could support (residual) adaptation mechanisms to recover a “healthy set point” on the longer run. Here, at least in principle, MVS could aid; in particular because the MVS-induced changes in spatial attention are not short-lived but consistently present for at least one hour of continuous stimulation, as we do show here in healthy subjects.

In summary, MVS at 3T produces sustained effects on the VOR and on spatial attention for periods of at least one hour and are accompanied by central adaptive changes, as evident from short-lived aftereffects. Our results do not only have important implications for the design and interpretation of fMRI studies. They also provide opportunities for basic research of the vestibular system and its influence on neural processing, behaviour and cognition. Finally, sustained MVS bears great clinical potential, counteracting pathological biases of attention and exploration in stroke patients.

## Data Availability

The datasets analysed during the current study are available from the corresponding author on reasonable request.

## Notes

### Competing Interest Statement

The authors have declared no competing interest.

## References

Boegle, R., Kirsch, V., Gerb, J., & Dieterich, M. (2020). Modulatory effects of magnetic vestibular stimulation on resting-state networks can be explained by subject-specific orientation of inner-ear anatomy in the MR static magnetic field. Journal of Neurology, 267(Suppl 1), 91–103. 10.1007/s00415-020-09957-3

Boegle, R., Ertl, M., Stephan, T., & Dieterich, M. (2017). Magnetic vestibular stimulation influences resting-state fluctuations and induces visual-vestibular biases. Journal of Neurology, 264(5), 999–1001. 10.1007/s00415-017-8447-6

Boegle, R., Stephan, T., Ertl, M., Glasauer, S., & Dieterich, M. (2016). Magnetic vestibular stimulation modulates default mode network fluctuations. NeuroImage, 127, 409–421. 10.1016/j.neuroimage.2015.11.065

Ertl, M., & Boegle, R. (2019). Investigating the vestibular system using modern imaging techniques-A review on the available stimulation and imaging methods. Journal of Neuroscience Methods, 326, 108363. 10.1016/j.jneumeth.2019.108363

Ferrè, E. R., & Haggard, P. (2020). Vestibular cognition: State-of-the-art and future directions. Cognitive Neuropsychology, 37(7-8), 413–420. 10.1080/02643294.2020.1736018

Go, C. C., Taskin, H. O., Ahmadi, S.-A., Frazzetta, G., Cutler, L., Malhotra, S., Morgan, J. I. W., Flanagin, V. L., & Aguirre, G. K. (2022). Persistent horizontal and vertical, MR-induced nystagmus in resting state Human Connectome Project data. NeuroImage, 255, 119170. 10.1016/j.neuroimage.2022.119170

Holé, J., Reilly, K. T., Nash, S., & Rode, G. (2020). Caloric Vestibular Stimulation Reduces the Directional Bias in Representational Neglect. Brain Sciences, 10(6). 10.3390/brainsci10060323

Jareonsettasin, P., Otero-Millan, J., Ward, B. K., Roberts, D. C., Schubert, M. C., & Zee, D. S. (2016). Multiple Time Courses of Vestibular Set-Point Adaptation Revealed by Sustained Magnetic Field Stimulation of the Labyrinth. Current Biology : CB, 26(10), 1359–1366. 10.1016/j.cub.2016.03.066

Karnath, H.-O., Fetter, M., & Dichgans, J. (1996). Ocular exploration of space as a function of neck proprioceptive and vestibular input - observations in normal subjects and patients with spatial neglect after parietal lesions. Experimental Brain Research(109), 333–342.

Karnath, H.-O. (1994b). Disturbed coordinate transformation in the neural representation of space as the crucial mechanism leading to neglect. Neuropsychological Rehabilitation(4(2)), 147–150.

Karnath, H.-O. (1994). Subjective body orientation in neglect and the interactive contribution of neck muscle proprioception and vestibular stimulation. Brain, 1001–1012.

Karnath, H.-O. (2015). Spatial attention systems in spatial neglect. Neuropsychologia, 75, 61–73. 10.1016/j.neuropsychologia.2015.05.019

Karnath, H.-O., & Rorden, C. (2012). The anatomy of spatial neglect. Neuropsychologia, 50(6), 1010–1017. 10.1016/j.neuropsychologia.2011.06.027

Karnath, H.-O., Rosenzopf, H., Smaczny, S., & Lindner, A. (2022). Spatial neglect after stroke is reduced when lying inside a 3T MRI scanner. BioRxiv. Advance online publication. 10.1101/2022.08.01.502290

Lindner, A., Wiesen, D., & Karnath, H.-O. (2021). Lying in a 3T MRI scanner induced neglect-like spatial attention bias. ELife. Advance online publication. 10.7554/eLife.71076

Mian, O. S., Glover, P. M., & Day, B. L. (2015). Reconciling Magnetically Induced Vertigo and Nystagmus. Frontiers in Neurology, 6, 201. 10.3389/fneur.2015.00201

Mian, O. S., Li, Y., Antunes, A., Glover, P. M., & Day, B. L. (2013). On the Vertigo Due to Static Magnetic Fields. PLoS ONE. Advance online publication. 10.1371/journal.pone.0078748.g001

Mian, O. S., Li, Y., Antunes, A., Glover, P. M., & Day, B. L. (2016). Effect of head pitch and roll orientations on magnetically induced vertigo. The Journal of Physiology, 594(4), 1051–1067. 10.1113/JP271513

Owens, A. D., & Leibowitz, H. W. (1980). Accommodation, Convergence, and Distance Perception in Low Illumination: 540-50. American Journal of Optometry and Physiological Optics(57(9)), 540–550.

Roberts, D. C., Marcelli, V., Gillen, J. S., Carey, J. P., Della Santina, C. C., & Zee, D. S. (2011). Mri magnetic field stimulates rotational sensors of the brain. Current Biology : CB, 21(19), 1635–1640. 10.1016/j.cub.2011.08.029

Rubens, A. B. (1985). Caloric stimulation and unilateral visual neglect. Neurology. Advance online publication. 10.1212/WNL.35.7.1019

Vallar, G., Papagno, C., Rusconi, M. L., & Bisiach, E. (1995). Vestibular stimulation, spatial hemineglect and dysphasia, selective effects. Cortex; a Journal Devoted to the Study of the Nervous System and Behavior, 31(3), 589–593. 10.1016/s0010-9452(13)80070-6

Ward, B. K., Otero-Millan, J., Jareonsettasin, P., Schubert, M. C., Roberts, D. C., & Zee, D. S. (2017). Magnetic Vestibular Stimulation (MVS) As a Technique for Understanding the Normal and Diseased Labyrinth. Frontiers in Neurology, 8, 122. 10.3389/fneur.2017.00122

Ward, B. K., Roberts, D. C., Otero-Millan, J., & Zee, D. S. (2019). A decade of magnetic vestibular stimulation: From serendipity to physics to the clinic. Journal of Neurophysiology, 121(6), 2013–2019. 10.1152/jn.00873.2018

Zee, D. S., Jareonsettasin, P., & Leigh, R. J. (2017). Ocular stability and set-point adaptation. Philosophical Transactions of the Royal Society of London. Series B, Biological Sciences, 372(1718). 10.1098/rstb.2016.0199

